# Spatial and temporal organization of RecA in the *Escherichia coli* DNA-damage response

**DOI:** 10.1101/413880

**Authors:** Harshad Ghodke, Bishnu P Paudel, Jacob S Lewis, Slobodan Jergic, Kamya Gopal, Zachary J Romero, Elizabeth A Wood, Roger Woodgate, Michael M Cox, Antoine M van Oijen

**Affiliations:** School of Chemistry, University of Wollongong, Wollongong, Australia; Illawarra Health and Medical Research Institute, Wollongong, Australia; Biochemistry Department, University of Wisconsin-Madison, Madison, Wisconsin, United States of America; Laboratory of Genomic Integrity, National Institute of Child Health and Human Development, National Institutes of Health, Bethesda, MD, USA

## Abstract

The RecA protein orchestrates the cellular response to DNA damage via its multiple roles in the bacterial SOS response. Lack of tools that provide unambiguous access to the various RecA states within the cell have prevented understanding of the spatial and temporal changes in RecA structure/function that underlie control of the damage response. Here, we develop a monomeric C-terminal fragment of the λ repressor as a novel fluorescent probe that specifically interacts with RecA filaments on single-stranded DNA (RecA*). Single-molecule imaging techniques in live cells demonstrate that RecA is largely sequestered in storage structures during normal metabolism. Upon DNA damage, the storage structures dissolve and the cytosolic pool of RecA rapidly nucleates to form early SOS-signaling complexes, maturing into DNA-bound RecA bundles at later time points. Both before and after SOS induction, RecA* largely appears at locations distal from replisomes. Upon completion of repair, RecA storage structures reform.

## Introduction

All cells possess an intricately regulated response to DNA damage. Bacteria have evolved an extensive regulatory network called the SOS response to control the synthesis of factors that protect and repair the genome. Processes coordinately regulated within the SOS response include error-free DNA repair, error-prone lesion bypass, cell division, and recombination.

The RecA protein is the master regulator of SOS, with at least three distinct roles. First, RecA forms a ternary complex with single-stranded DNA (ssDNA) and ATP to form the activated RecA*. RecA* catalyzes auto-proteolysis of the transcriptional repressor LexA to induce expression of more than 40 SOS genes (Courcelle et al., 2001; Fernandez De Henestrosa et al., 2000; Kenyon and Walker, 1980; Little and Mount, 1982; Little et al., 1981). RecA* thus uses the ssDNA generated when replication forks encounter DNA lesions as an induction signal (Sassanfar and Roberts, 1990). Second, along with several other accessory proteins, RecA mediates error-free recombinational DNA repair at sites of single-strand gaps, double-strand breaks (DSBs) and failed replisomes (Cox et al., 2000; Kowalczykowski, 2000; Kuzminov, 1995; Lusetti and Cox, 2002). Third, the formation and activity of active DNA Polymerase V complex capable of lesion bypass requires RecA* (Jaszczur et al., 2016; Jiang et al., 2009; Robinson et al., 2015).

RecA is a prototypical member of a class of proteins that are critical for genomic stability across all domains of life (Baumann et al., 1996; Bianco et al., 1998; Lusetti et al., 2003a; San Filippo et al., 2008; Sung, 1994). In higher organisms, including humans, the homologous protein Rad51 supports error-free double-strand break repair by catalyzing strand exchange much like the RecA protein does in eubacteria (Baumann et al., 1996; Sung, 1994). Mutations in human Rad51 and accessory proteins have been implicated in carcinomas and Fanconi anemia (Chen et al., 2015; Kato et al., 2000; Prakash et al., 2015). Unsurprisingly, RecA and related recombinases are highly regulated, with a variety of accessory proteins governing every facet of their multiple functions (Cox, 2007). Directed-evolution approaches can be used to enhance the catalytic activities of recombinases in cells (Kim et al., 2015). However, RecA functional enhancement has a cost, disrupting an evolved balance between the various processes of DNA metabolism that share a common genomic DNA substrate (Kim et al., 2015). Many deleterious genomic events occur at the interfaces between replication, repair, recombination, and transcription.

An understanding of how organisms maintain genetic integrity requires an examination of the protein actors in their native cellular environments. In response to DNA damage, transcription of the *recA* gene is upregulated ten-fold within minutes (Courcelle et al., 2001; Renzette et al., 2005). Using immunostaining, the copy number of RecA in undamaged cells has been estimated to be about 7,000-15,000 per cell, increasing to 100,000 per cell upon triggering the DNA-damage response (Boudsocq et al., 1997; Stohl et al., 2003). Visualization of C-terminal GFP fusions of wild-type and mutant *recA* alleles placed under the native *recA* promoter in *E. coli* have revealed that RecA forms foci in cells (Lesterlin et al., 2014; Renzette et al., 2005; Renzette et al., 2007). Interpretation of the localizations observed in these experiments has been clouded by three issues: 1. fluorescent fusions of RecA have consistently yielded loss-of-function phenotypes to RecA (Handa et al., 2009; Renzette et al., 2005), making interpretation of the localizations revealed by these tagged proteins highly challenging. 2. this issue is further complicated by the fact that fluorescent proteins do not behave as inert tags and can influence intracellular localization in bacterial cells (Ghodke et al., 2016; Ouzounov et al., 2016). Indeed, *Bacillus subtilis* RecA tagged with GFP, YFP and mRFP yielded different localizations in response to DNA damage (Kidane and Graumann, 2005). These challenges do not come as a surprise since both N- and C-terminal ends are important for RecA function and localization (Eggler et al., 2003; Lusetti et al., 2003a; Lusetti et al., 2003b; Rajendram et al., 2015). 3. at least *in vitro*, untagged RecA has a remarkable ability to self-assemble, into different complexes that form on single-stranded DNA (RecA*), on double-stranded DNA, or are free of DNA (Brenner et al., 1988; Egelman and Stasiak, 1986, 1988; Stasiak and Egelman, 1986). The properties of these assemblies are often determined by the state of hydrolysis of associated ATP. Thus, unambiguous assignment of the molecular composition of RecA features in live cells has been difficult.

In the absence of DNA, RecA can polymerize to form aggregates of various stoichiometry to yield dimers, tetramers, ‘rods’ and ‘bundles’ (Brenner et al., 1988). Some of these states may have a physiological relevance: RecA fusions with the best functionality have revealed DNA-free aggregates that are confined to the cellular poles, outside of the nucleoid and associated with anionic phospholipids in the inner membrane (Rajendram et al., 2015; Renzette et al., 2005). These DNA-free aggregates were hypothesized to be ‘storage structures’ of RecA, although their functionality in and relevance to the DNA damage response remain unclear.

Early electron-microscopy studies revealed that multiple dsDNA-RecA-ATPγS filaments could also associate to form structures confusingly termed as ‘bundles’ (Egelman and Stasiak, 1988). This study also identified that ssDNA-RecA-ATPγS filaments could aggregate together. Electron microscopy of cells revealed that RecA appeared to form ‘bundles’ that were aligned next to the inner membrane in cells after DNA damage (Levin-Zaidman et al., 2000). In cells carrying an additional allele of wild-type RecA at a secondary chromosomal locus to increase overall RecA function, long RecA structures called ‘bundles’ were formed during double-strand break repair (Lesterlin et al., 2014). These bundles are similar to RecA structures called ‘threads’, that nucleate at engineered double-strand breaks in *Bacillus subtilis* (Kidane and Graumann, 2005). RecA bundles form after SOS induction by other means than double-strand breaks, and also then interact with anionic phospholipids in the inner membrane (Garvey et al., 1985; Rajendram et al., 2015). The appearance of elongated RecA* foci after treatment with UV has not always been associated with bundle formation (Renzette et al., 2007). It should be noted that whereas assemblies of RecA observed *in vivo* have been variously referred to as filaments, threads or bundles, their correspondence to the *in vitro* observations of RecA aggregates referred to as ‘rods’ or ‘bundles’ remains unclear.

Due to the similar morphology of the fluorescence signal arising from these various DNA-bound repair or DNA-free storage structures, teasing out dynamics of individual repair complexes in live cells has proven difficult. The limited functionality of RecA fusion proteins utilized to date also raises concerns about the relationship of the observed structures to normal RecA function. Several fundamental questions remain unanswered: When and where does SOS signaling occur in cells? How is excess RecA stored?

In this work, we describe the development of a probe that specifically visualizes RecA structures on DNA, and utilize it as part of a broader effort to provide a detailed time line of RecA structural organization in living cells after DNA damage. With the objective of selectively localizing DNA-bound and ATP-activated RecA* as a key repair intermediate inside living cells, we produced a <u>m</u>onomeric, catalytically dead N-terminal truncation of the bacteriophage λ repressor <u>CI</u> (mCI*; CI 101-229, A152T, P158T, K192A*) that retains the ability to bind RecA-ssDNA filaments. Removal of the N-terminal domain renders the mCI unable to bind DNA, leaving only RecA* as a binding partner. Using both untagged and fluorescently labelled mCI constructs, we document the effects of mCI *in vitro* and *in vivo*. We use mCI as well as the most functional RecA-GFP fusion protein variants to distinguish between the various types of RecA structures and follow their behavior through time. In addition, we examine the location of RecA* foci formed in the nucleoid in relation to the location of the cellular replisomes. Our results reveal how the activity of RecA is regulated upon triggering of the SOS pathway and identify the various states of RecA that are relevant throughout the damage response.

## Results

### RecA-GFP forms different types of aggregates in cells

Damage in the template strand can result in the stalling or decoupling of replication, leading to the accumulation of single-stranded DNA. This ssDNA provides a RecA nucleation site to form RecA*, the structure that amplifies the cellular signal for genetic instability (Figure 1A). In response to DNA damage, transcription of the SOS-inducible genes is up-regulated. Because production of RecA occurs rapidly after damage, it is critical to observe live cells at early time points with high temporal resolution after SOS induction.

**Figure 1:**
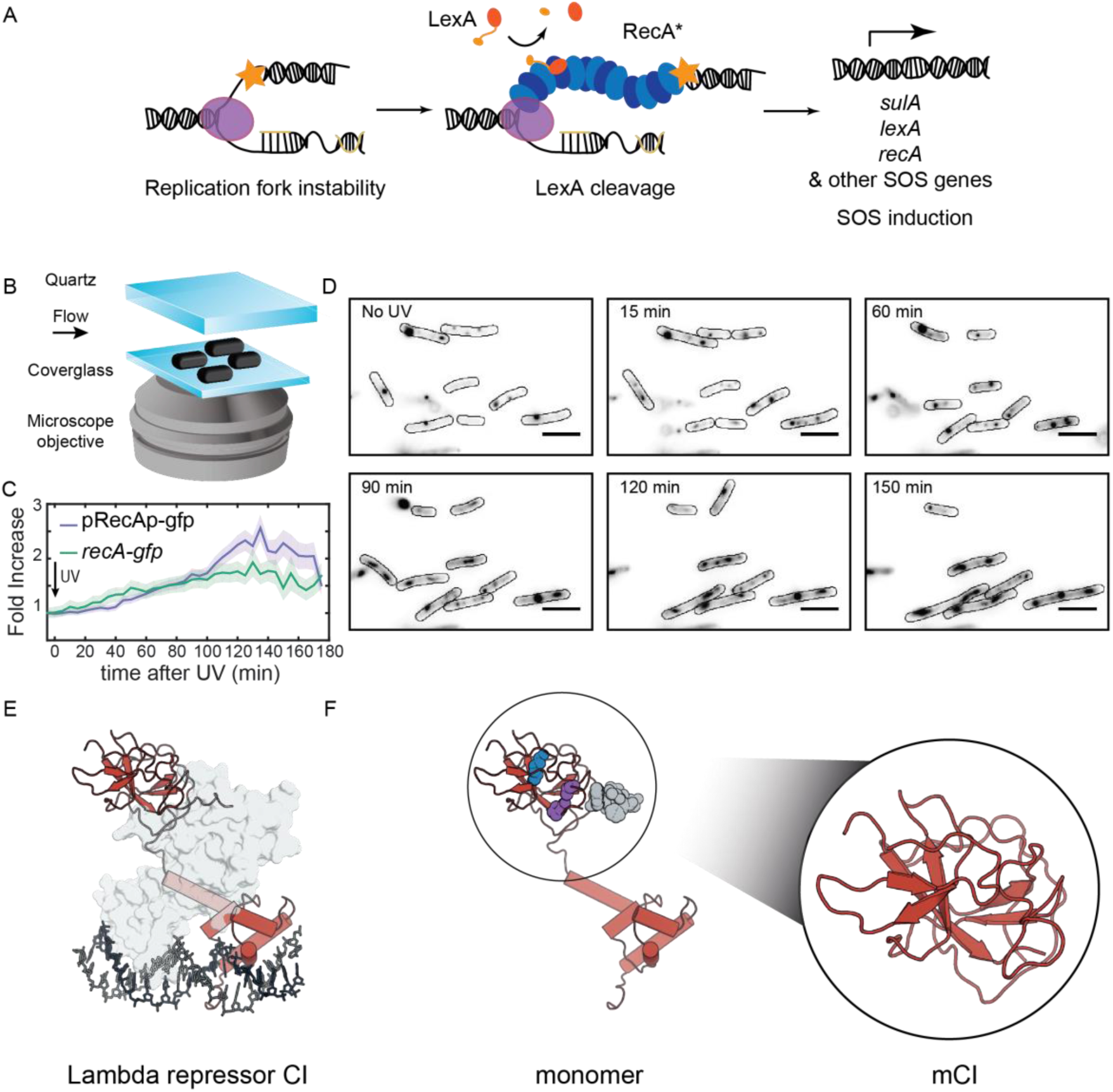
RecA forms different intracellular structures in response to UV irradiation. **A.** Consensus model for SOS induction after DNA damage, illustrating the formation of ssDNA-containing RecA* filaments at sites of stalled replication forks. These RecA* filaments induce the SOS response by promoting cleavage of LexA. **B.** Schematic of flow-cell setup for live-cell imaging. **C.** Plots of relative increase in mean intensity of GFP in pRecAp-gfp *cells* (purple, strain# HG260) or RecA-GFP expressed from the native chromosomal locus (*recA-gfp* cells). Cells are irradiated with 20 Jm^−2^ of UV at *t* = 0 min. Shaded error bars represent standard error of the mean cellular fluorescence measured in cells. 50 – 200 cells were analyzed at each time point. Scale bar corresponds to 5 μm. See also SI Movie 1. **D.** Imaging of *recA-gfp* cells (strain# HG195) reveals that RecA-GFP forms foci of various morphologies at different stages during the SOS response. Stills from SI Movie 2 are presented here. **E.** Crystal structure of the operator bound dimeric λ repressor CI (PDB ID: 3BDN). **F.** Monomer of CI showing the catalytic lysine (K192, purple), residues that mediate dimerization (A152 and P158, blue), and the C terminus involved in dimerization (grey). Inset shows the monomeric C-terminal fragment ‘mCI’ defined as CI(101-229, A152T P158A and K192A) used in this study.

With the objective of characterizing the spatial and temporal organization of RecA in cells during SOS induction, we performed time-lapse imaging of individual *E. coli* cells immobilized in flow cells using a variety of fluorescent probes (See SI for imaging methods). This setup enabled us to monitor growing cells with nutrient flow at 30°C, while keeping the cells in place to support long-term, time-lapse imaging of individual cells. A quartz window in the flow cell enabled us to provide *in situ* UV irradiation with a defined dose (20 Jm^−2^) at the start of the experiment. Following this, we monitored fluorescence every 5 minutes over the course of three hours by wide-field acquisition (Figure 1B). For this study we chose to induce SOS with UV for two key reasons: first, UV light is a strong inducer of the SOS response, and second, a pulse of UV light serves to synchronize the DNA damage response in cells that are continuously replicating DNA without the need for additional synchronization.

First, we set out to characterize the temporal activity of the *recA* promoter alone in wild-type MG1655 cells in response to ultraviolet radiation. We imaged cells that express fast-folding GFP from the *gfpmut2* gene placed under the *recA* promoter on a low-copy reporter plasmid, with a maturation time of less than 5 minutes (‘pRecAp-gfp’ cells; strain# HG260) (Kalir et al., 2001; Zaslaver et al., 2006). The copy number of this reporter plasmid has been shown to remain unchanged in response to ultraviolet radiation (Ronen et al., 2002). The cells retain the chromosomal copy of the wild type *recA* gene. Measurements of the mean fluorescence intensity of cytosolic GFP in pRecAp-gfp cells exhibited a gradual increase peaking at approximately 135 min, with a maximum that corresponded to a two-fold increase compared to the initial fluorescence intensity (SI Movie 1 & Figure 1C). After UV exposure, the accumulation of the fluorescent reporter protein expressed from the *recA* promoter reaches a maximum only after more than two hours. By extension, this gradual increase is used here to define the time period during which cellular RecA concentration is increasing after UV treatment.

Next, we imaged MG1655 cells that carry a *recA-gfp* fusion allele expressed from the *recAo1403* operator in place of the wild-type chromosomal copy of *recA* (‘*recA-gfp*’ cells; strain# HG195). The *recAo1403* promoter increases the basal (non-SOS) level of *recA* expression by a factor of 2-3 (Rajendram et al., 2015; Renzette et al., 2005). Despite the higher expression level, cells expressing this RecA-GFP fusion protein are deficient in RecA functions; notably, these cells exhibit a three-fold lower survival in response to UV irradiation, and ten-fold lower ability to perform recombination (Renzette et al., 2005). Additionally, these cells exhibit delayed kinetics of SOS induction, but are still able to induce the SOS response to the same extent as wild-type cells (Renzette et al., 2005). In response to UV irradiation, GFP fluorescence in *recA-gfp* cells increased after DNA damage and peaked at approximately 130 min (SI Movie2 & Figure 1C & 1D). Thus, the kinetics of the observed increase in the levels of chromosomally expressed RecA-GFP fusion protein are the same those of the increase seen with the plasmid-based *gfpmut2* reporter under control of the *recA* promoter.

Measurements of the abundance of the *recA* transcript after SOS induction have revealed a ten-fold increase within minutes after irradiation with UV (Courcelle et al., 2001). In bulk experiments, the amount of RecA protein has been shown to attain a maximum at 90 min after introduction of damage (Salles and Paoletti, 1983). Results from our live-cell experiments are generally consistent with these studies, revealing that the amount of RecA accumulated in cells attains a maximum at a time after triggering the SOS response that is much later than the de-repression of the *recA* promoter. During the SOS response, many cells undergo filamentation as cell division is blocked while some DNA synthesis continues (Howard-Flanders et al., 1968). The increase in gene expression counters the dilution in the cellular RecA concentration that is caused by the filamentation of the cells.

In these experiments, RecA-GFP formed well-defined features both before and after DNA damage (SI Movie 2 and Figure 1D). These foci exhibited various morphologies ranging from punctate foci to bundles. The foci became generally larger and more abundant after UV irradiation. To determine whether RecA foci formed in the absence of DNA damage are functionally distinct from those formed during the SOS response, we set out to specifically visualize RecA*, the complex that is formed when RecA binds ssDNA and that is actively participating in repair. To that end, we investigated interaction partners of the ssDNA-RecA filament that are not endogenously present in *E. coli*. Since the MG1655 strain we use in our studies is cured of bacteriophage λ, we focused on co-opting the λ repressor to detect RecA* in cells (Figure 1E) (Roberts and Roberts, 1975).

### The monomeric C-terminal fragment of the bacteriophage λ repressor (mCI) stabilizes dynamics of RecA-ssDNA filaments *in vitro*

The bacteriophage λ repressor CI is responsible for the maintenance of lysogeny in *E. coli* infected with phage λ (Echols and Green, 1971). Oligomers of CI bind the operator regions in the constitutive P_L_ and P_R_ promoters in λ DNA and inhibit transcription from these promoters (Ptashne et al., 1980). In response to DNA damage, the λ repressor CI exhibits RecA* dependent auto-proteolysis, much like the homologous proteins in bacteria, LexA and UmuD (Burckhardt et al., 1988; Ferentz et al., 1997; Luo et al., 2001; Roberts and Roberts, 1975; Stayrook et al., 2008; Walker, 2001). In this reaction, the ssDNA-RecA filament (RecA*) stabilizes a proteolysis-competent conformation of CI enabling auto-proteolysis at Ala111-Gly112 (Ndjonka and Bell, 2006; Sauer et al., 1982). This co-protease activity of the RecA* filament results in loss of lysogeny due to de-repression of transcription of *cI* and prophage induction of λ. The N-terminal DNA-binding domain of CI is dispensable for interactions with RecA*(Gimble and Sauer, 1989). A minimal C-terminal fragment of the λ repressor CI(101-229 A152T P158T K192A) (henceforth referred to as mCI, Molecular weight 14307.23 Da; Figure 1F) efficiently inhibits the auto-catalytic cleavage of a hyper-cleavable monomeric C-terminal fragment CI(92-229) (Ndjonka and Bell, 2006). Cryo-electron microscopy has revealed that the mCI binds deep in the groove of the RecA filament (Galkin et al., 2009).

Given the existing extensive *in vitro* characterization of mCI, we decided to further develop it as a probe for detecting RecA* in cells. To better understand the kinetics, cooperativity and affinity of mCI for RecA-ssDNA filaments, we first pursued an *in vitro* investigation of the interaction between mCI and RecA filaments. Additionally, to visualize mCI in live-cell experiments, we made fusion constructs with fluorescent proteins tagged to the N terminus of mCI via a 14-amino acid linker. To perform time-lapse imaging, we tagged mCI with the yellow fluorescent protein YPet, and to perform live-cell photoactivatable light microscopy (PALM), we tagged mCI with the photoactivatable red fluorescent protein PAmCherry. Untagged mCI and the two fluorescently labelled constructs, PAmCherry-mCI and YPet-mCI were purified and characterized for RecA-ssDNA binding as described below (See SI for details, SI Figure 1A).

Binding of the mCI constructs to ssDNA-RecA filaments was first assayed by surface plasmon resonance (SPR). We immobilized a 5’ biotinylated (dT)_71_ ssDNA substrate on the surface of a streptavidin-functionalized SPR chip (Figure 2A) and assembled RecA-ssDNA filaments by injecting 1 μM RecA in buffer supplemented with ATP. This was followed by injection of buffer without RecA, but supplemented with ATPγS to minimize disassembly of the RecA filament on the ssDNA immobilized on the chip surface (SI Figure 1B). Next, the experiment was repeated but now introducing to pre-formed RecA* filaments solutions that not only contain stabilizing ATPγS, but also either untagged or fluorescently tagged mCI proteins. Figure 2B shows scaled sensorgrams that are corrected for any disassembly of the ssDNA-RecA-ATPγS filament and thus reporting on interactions of mCI with the highly stable RecA* filament (see also SI Figure 1C). These sensorgrams reveal that mCI associates with the RecA filament in a biphasic manner. Dissociation of mCI from the RecA filament was slow, with a dissociation halftime (t_1/2_) of 850 s. In comparison, the fluorescently tagged constructs dissociated faster, but still slowly enough for use as a probe for the detection of RecA*. We measured a t_1/2_ = 260 s and 280 s for YPet-mCI and PAmCherry-mCI respectively.

**Figure 2:**
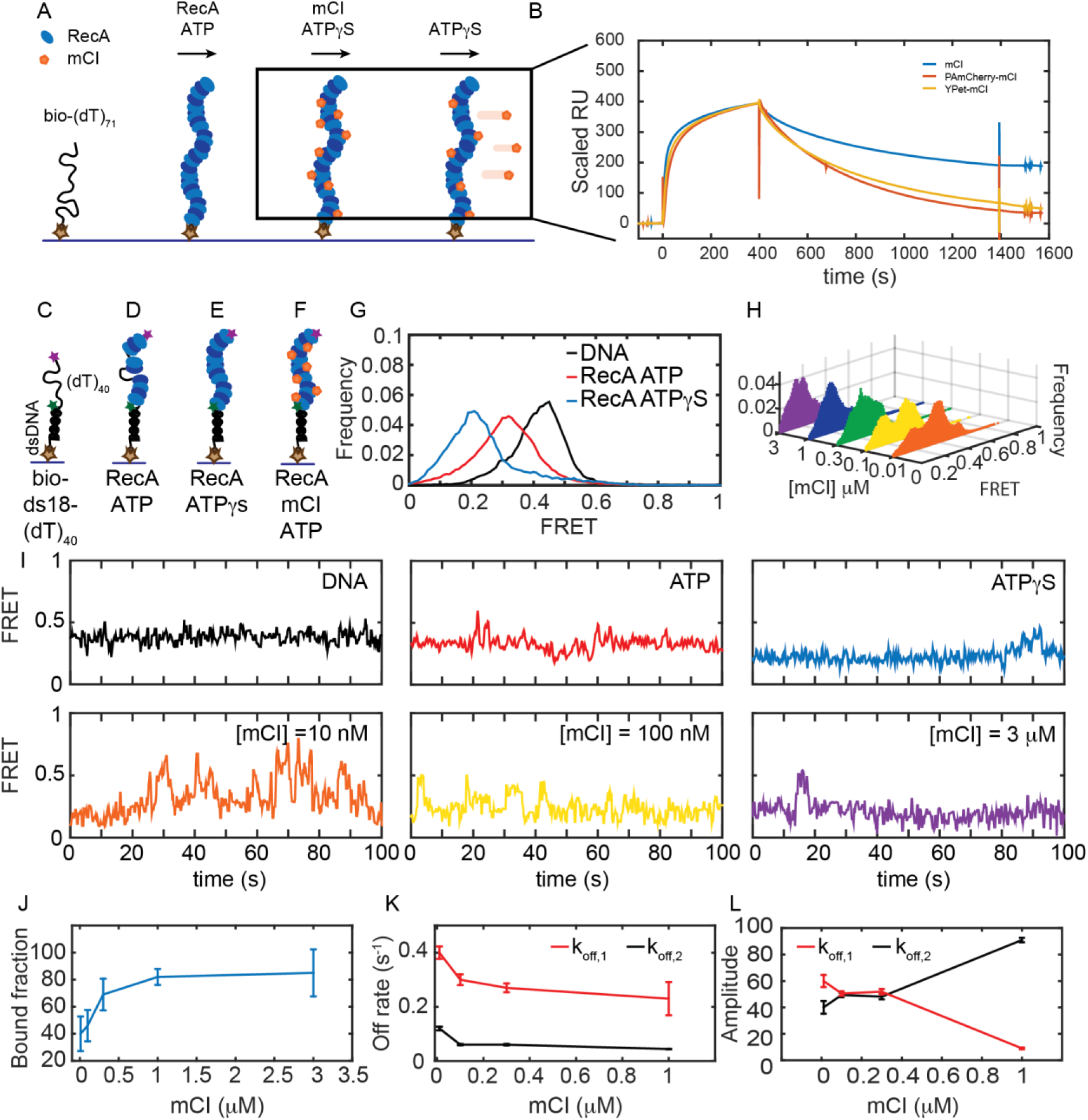
mCI stabilizes ssDNA-RecA filaments *in vitro*. **A.** Schematic of SPR experiment probing association and dissociation kinetics of mCI from ssDNA-RecA-ATPγS filaments on the surface of an SPR chip. ssDNA-RecA-ATPγS filaments were assembled on a biotinylated (dT)_71_ ssDNA molecule. mCI, YPet-mCI or PAmCherry-mCI were then flowed into the flow cell at time *t* = 0 for 400 s to monitor the association phase. Dissociation of mCI from ssDNA-RecA-ATPγS filaments was observed by leaving out mCI from the injection buffer. **B.** Sensorgram reveals biphasic association of mCI to RecA filaments, followed by a slow dissociation from the ssDNA-RecA-ATPγS filament. Sensorgrams presented here are corrected for slow disassembly of the RecA-ATPγS filament, and data are scaled to the binding curve of YPet-mCI for purposes of illustration (see also SI Figure 1 for unscaled data). **C.** Schematic of single-molecule FRET assay used to probe the influence of mCI binding on the conformational state of the ssDNA-RecA-ATP filament assembled on a ssDNA (dT)_40_ overhang. Biotinylated substrate DNA (bio-ds18-(dT)_40_) was immobilized on a functionalized coverslip via a streptavidin-biotin interaction. **D.** RecA binds the ssDNA overhang dynamically to form a ssDNA-RecA filament. **E.** In the presence of ATPγS, RecA forms a stable filament. **F.** Incubation with mCI leads to a RecA filament decorated with mCI. **G.** FRET distributions observed from the substrate alone, with RecA-ATP and RecA-ATPγS. **H.** Titration of mCI shifts the RecA-ATP distribution to that of the active filament. **I.** Example FRET traces of DNA substrate alone or when bound to RecA in the presence of ATPγS, or when bound to RecA in the presence of ATP and mCI (0, 10, 100, 300, 1000 and 3000 nM mCI) **J.** Fitting of the Hill equation to the percentage of bound fraction as a function [mCI] reveals a *K*_*D*_ of 36 ± 10 nM and a cooperativity of 2.4 ± 0.2 **K.** Off rates measured from binding of mCI to ssDNA-RecA-ATP filaments. **L.** Percentage amplitude of the detected rate-constants as a function of [mCI] reveals enrichment of the population decaying according to the slow off-rate as a function of [mCI]. See also SI Figures 1 and 2.

To probe the influence of mCI binding on the conformational state of the RecA* filament, we next studied association of mCI with ssDNA-RecA-ATP filaments using single-molecule Förster Resonance Energy Transfer (FRET) experiments. We used a previously described DNA substrate consisting of a biotinylated 18-mer double-stranded region preceded by a 5’ (dT)_40_ overhang (‘bio-ds18-(dT)_40_’, See SI for details, Figure 2C) (Park et al., 2010). This substrate represents the partly single-stranded and partly double-stranded nature of the DNA substrate that is thought to be generated in the context of replisomes encountering lesions *in vivo*. The ssDNA region is labelled with a Cy3 donor probe on one end and a Cy5 acceptor probe on the other so that the degree of extension of the ssDNA can be measured by FRET. The DNA substrate was immobilized on a streptavidin-coated surface in a flow cell and the Cy5 FRET signal was measured upon excitation of the Cy3 dye with a 532-nm laser (see SI for details). Consistent with previous FRET investigations of this DNA substrate (Park et al., 2010), the DNA substrate alone exhibited a FRET distribution with a mean value of 0.43 ± 0.07 (mean ± standard deviation of a single Gaussian fit to the data) reflecting the ability of the ssDNA overhang to entropically collapse and sample a large number of conformational states (Figure 2C, 2G & 2I; see ‘DNA’ trace). In the presence of ATP and RecA, the resulting FRET distribution exhibited a peak with a mean FRET value of 0.3 ± 0.1, consistent with the formation of a highly dynamic RecA filament undergoing simultaneous assembly and disassembly (Figure 2D, 2G and 2I ‘ATP’ trace). Upon incubating the DNA substrate with RecA in the presence of ATPγS, we observed a shift in the FRET distribution to an even lower value of 0.2 ± 0.07, reflecting the formation of a rigid, fully extended ssDNA-RecA filament (Figure 2E, 2G and 2I ‘ATPγS’). Since ATPγS traps the RecA filament in an ‘active’ conformation that is capable of LexA repressor autocatalytic cleavage, we interpreted the 0.2 FRET state as corresponding to the active state (Craig and Roberts, 1981). Incubation of RecA with ADP revealed a broad FRET distribution similar to that obtained in the presence of ATP, reflecting unstable RecA filaments assembled on the DNA overhang (See SI Figure 2A).

Next, we studied the FRET displayed by the ssDNA-RecA-ATP filament while titrating in purified mCI (Figure 2H and SI Figure 2B) to gain insight into the influence of mCI binding on the stability of ssDNA-RecA-ATP filaments (Figure 2F). In the presence of mCI the FRET substrate exhibited a bi-modal behavior: either molecules exhibited the 0.43 FRET state or the 0.2 FRET state. Upon increasing mCI concentration, the FRET distribution gradually shifted from a mean of 0.43 to 0.2 in response to higher concentrations of mCI (Figure 2H). By fitting the distributions to a sum of two Gaussian fits reflecting the ‘bound’ state (0.2 FRET) and ‘unbound’ state (0.4 FRET), we were able to obtain the bound fraction at every concentration of mCI tested (Figure 2H and 2J). Fitting these data to the Hill equation yielded an equilibrium dissociation constant of 36 ± 10 nM with a Hill coefficient of 2.4 ± 0.2 (Figure 2J). The increase in the population of molecules in the lowest FRET state in response to an increase in mCI concentration demonstrates that mCI stabilizes the RecA filament in the active conformation.

Examination of the FRET traces revealed that in the presence of mCI, the DNA substrate exhibits stochastic transitions from the RecA-bound to the unbound state (e.g., Figure 2I for [mCI] = 10 nM). The frequency of these transitions to the unbound state decreased in the presence of high concentration of mCI (Figure 2I, see also SI Figure 2B). FRET traces of DNA substrates in the presence of RecA and saturating concentrations of mCI (3 μM) exhibited stable, long-lived binding events at a FRET value of 0.2 over the time scale of acquisition (Figure 2G). To obtain off rates from the data, we applied a threshold of 0.3 (SI Figure 2C) to segment the trajectories such that segments with FRET values less than 0.3 were considered to reflect the ‘bound’ state, and those above 0.3 were considered to be the ‘unbound’ state. The cumulative residence time distributions for the binding events (low FRET values) in the FRET trajectories were best fit by a sum of two exponentials decaying according to a fast off rate k_off,1_ = 0.23 ± 0.06 s^−1^ and a slow off rate k_off,2_ = 0.044 ± 0.002 s^−1^ (Figure 2K). These off-rates were largely independent of the concentration of mCI (Figure 2K). However, strikingly, the fraction of the population decaying following the slower off rate increased from 35% in the absence of mCI to 91% in the presence of 1μM mCI (Figure2L).

Next we probed the influence of mCI on the ability of RecA* to perform three key catalytic functions *in vitro*: LexA cleavage, ATP hydrolysis and strand-exchange. Incubation of pre-formed RecA* filaments on circular ssDNA M13mp18 substrates with micromolar concentrations of mCI revealed a pronounced inhibition of RecA* ATPase activity (SI Figure 2D). The tagged mCI variants did not significantly inhibit RecA* ATPase activity at concentrations under 500 nM (SI Figure 2D). Monitoring of LexA cleavage by RecA* in the presence of mCI revealed that even high concentrations of mCI did not inhibit LexA cleavage, at best, mCI at micromolar concentrations slowed the kinetics of LexA cleavage (Supplementary Fig. 2E). Finally, whereas strand-exchange by RecA* was potently inhibited at micromolar concentrations of mCI (Supplementary Fig. 2F), tagged mCI constructs did not significantly inhibit strand-exchange activity at concentrations below 500 nM (Supplementary Fig. 2F).

Taken together, these *in vitro* investigations provide insights into the consequences of mCI binding on the activity of RecA*. We found that mCI stabilizes the RecA* filament in the ‘active’ conformation that is capable of LexA cleavage. At high concentrations (5 – 10 μM), mCI can inhibit ATP hydrolysis and strand-exchange by RecA*, and delay LexA cleavage. This is consistent with mCI binding to the RecA nucleoprotein filament groove as anticipated. Importantly, at low concentrations (10 - 100 nM) similar to those we eventually employed as a standard in vivo (as described below), these key activities of RecA* are not significantly affected by the presence of mCI or tagged variant. These findings emphasize the suitability of the use of mCI derived probes for probing RecA* function.

### mCI inhibits SOS induction in a concentration-dependent manner

We probed whether mCI interacts with ssDNA-RecA filaments (RecA*) in cells upon DNA damage and potentially inhibits the SOS response. To that end, we created live-cell imaging vectors that express either mCI or the PAmCherry-mCI fusion from the *araBAD* promoter in a tunable manner depending on the amount of L-arabinose provided in the growth medium (Guzman et al., 1995). The ability of cells to induce SOS was assayed using a previously described set of SOS-reporter plasmids that express GFP in response to DNA damage (Zaslaver et al., 2006). In this assay, we measured the fluorescence of fast-folding GFP expressed from the *gfpmut2* gene under the SOS-inducible *sulA* promoter on a low-copy plasmid (‘*sulAp-gfp*’)(Zaslaver et al., 2006). As a control, we also measured GFP fluorescence from the promoter-less parent vector (‘*gfp*’). The copy number of these SOS-reporter plasmids is not influenced by the ultraviolet radiation (Ronen et al., 2002). To measure the ability of mCI to inhibit the SOS response in cells, we co-transformed wild-type MG1655 cells with either the pBAD-mCI vector (‘*mcI*’), pBAD-PAmCherry-mCI vector (‘*PAmCherry-mcI*’) or an empty pBAD vector (‘pBAD’), and *sulA* reporter (‘*sulAp-gfp*’) or promoter-less vector (‘*gfp*’) to generate four strains: 1. cells that carry the empty pBAD vector and the promoter-less *gfp* vector (‘*gfp*+pBAD’, strain# HG257) 2. cells that carry the empty pBAD vector and the *sulA* reporter plasmid (‘*sulAp-gfp* + pBAD’, strain# HG258) 3. cells that carry the pBAD-mCI vector and the *sulA* reporter plasmid (‘*sulAp-gfp* + *mcI*’, strain# HG253) and 4. cells that carry the pBAD-PAmCherry-mCI vector and the SOS-reporter plasmid (‘*sulAp-gfp + PAmCherry-mcI*’, strain# HG285).

We then acquired time-lapse movies of these cells to observe the evolution of the SOS response over three hours after UV damage (Figure 3A). As expected, when cells carrying the *sulA* reporter plasmid and the empty pBAD vector (‘*sulAp-gfp* + pBAD’) were irradiated with a 20 Jm^−2^ dose of UV, we observed a robust increase in GFP fluorescence (Figure 3A; strain# HG258). In contrast, cells carrying the promoter-less control vector and the empty pBAD vector (‘*gfp*+pBAD’) vectors did not exhibit any increase in GFP fluorescence in response to UV (Figure 3A, summarized in Figure 3C; strain# HG257).

**Figure 3:**
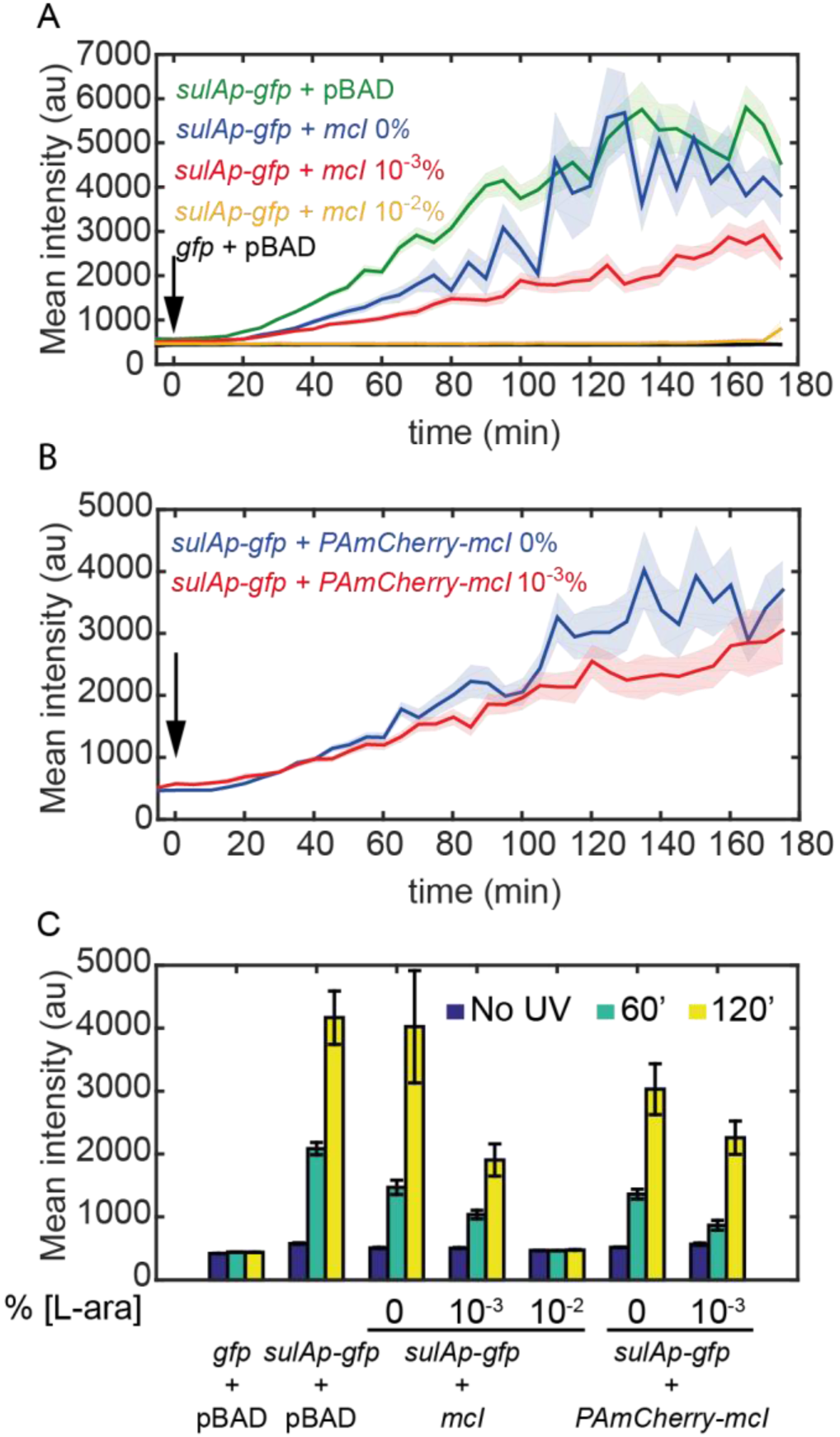
mCI inhibits the SOS response in a concentration-dependent manner. **A**. Time-lapse experiments were performed on MG1655 cells carrying the SOS-reporter plasmids (‘*gfp*’ or ‘*sulAp-gfp*’) and pBAD-mCI plasmid (‘*mcI’*) following irradiation with 20 Jm^-2^ of UV-irradiation at time *t* = 0 min. Mean intensity of GFP fluorescence was measured in cells carrying the reporter plasmid and mCI or empty vector, and plotted here as follows: ‘*sulAp-gfp* + pBAD’ cells (green; strain# HG258), ‘*gfp* + pBAD’ cells (black; strain# HG257), ‘*sulAp-gfp* + *mcI*’ (strain# HG253) (0% L-ara) (blue), 10^−3^ % L-ara (red) and 10^−2^ % L-ara (yellow), respectively. **B.** Mean intensity of GFP fluorescence in cells carrying the reporter plasmid and pBAD-PAmCherry-mCI plasmid (‘*sulAp-gfp* + *PAmCherry-mcI*’) (strain# HG285; 0% L-ara (blue) and 10^−3^ % L-ara (red)) is plotted as a function of time. Shaded error bars indicate standard error of mean cellular fluorescence for all cells imaged at the indicated time point. In these experiments, 50-200 cells were analyzed for each experiment for each of the 37 time points. **C.** Bar plots summarizing data presented in B and C under the indicated conditions at a time point before UV irradiation, one at 60, and one at 120 minutes after UV.

After these experiments confirming the robustness of the *sulA* reporter as a readout for SOS induction, we grew cells carrying both the *sulA* reporter and the mCI vectors (‘*sulAp-gfp* + *mcI*’, strain# HG253) in imaging medium containing 0, 10^−3^ or 10^−2^ % L-arabinose and immobilized them in flow cells. In this L-arabinose concentration regime, we expect the mCI copy number to be approximately 20, 50 and 500 copies per cell, respectively (Ghodke et al., 2016). Time-lapse acquisition after UV irradiation revealed that SOS induction was sensitive to the presence of mCI. Even leaky expression of mCI caused a measurable delay in GFP fluorescence (Figure 3A). This delay was found to be proportional to the expression level of mCI, and cells grown in 10^−2^ % L-arabinose exhibited nearly complete inhibition of SOS induction during the experimental timeline of three hours after UV irradiation (Figure 3A and 3C). Time-lapsed imaging of wild-type cells carrying the pBAD-PAmCherry-mCI vector and the *sulA* reporter plasmid (‘*sulAp-gfp* + *PAmCherry-mcI*’, strain# HG285) also revealed a similar delay in SOS induction depending on the concentration of L-arabinose in the growth medium (Figure 3B). These data suggest that mCI competes with LexA in cells at sites of RecA* in response to DNA damage.

### Most RecA filaments are formed at sites distal to replisomes after DNA damage

A long-standing model for SOS induction predicts that RecA* filaments are formed on chromosomal DNA when replisomes encounter UV lesions (Sassanfar and Roberts, 1990). These RecA* filaments are believed to be the sites of SOS induction. While several lines of evidence support the model that RecA* filaments are formed after UV irradiation, direct visualization in living *E. coli* cells has not been demonstrated. Given that mCI robustly interacts with RecA* filaments, we set out to visualize RecA* in cells exposed to UV light. Importantly, considering that mCI can inhibit RecA* activities at μM concentrations, we chose to express mCI variants from the tightly repressed, and tunable pBAD promoter. Incubating cells with low concentrations of L-arabinose results in cells expressing tagged mCI constructs in the 10-100 nM range in cells.

This was achieved by imaging a two-color strain that expresses a chromosomal YPet fusion of the *dnaQ* gene (that encodes the replisomal protein ϵ, a subunit of the replicative DNA polymerase III), and PAmCherry-mCI from the pBAD-PAmCherry-mCI plasmid in the presence of L-arabinose (strain# HG267). The YPet fusion has previously been shown to minimally affect the function of ϵ (Reyes-Lamothe et al., 2010; Robinson et al., 2015). The two-color strain was grown in medium containing small amounts of L-arabinose (5×10^−4^ %) to induce low expression of PAmCherry-mCI. Under these conditions, UV-surivival of HG267 was indistinguishable from that of MG1655/pBAD-mycHisB(strain# HG116) (SI Figure 3A). This was followed by time-lapse imaging (5 min intervals for 3 h after 20 Jm^−2^ of UV) in flow-cells. In this experiment, we performed live-cell PALM to detect PAmCherry-mCI, and TIRF imaging to detect replisomes (see SI for technical details related to imaging, SI Figure 3B). This approach enabled us to visualize replisomes as well as mCI foci (Figure 4A). To compare cells in the presence and absence of SOS induction, we carried out the same experiments in cells carrying the *lexA3*(Ind^-^) allele (strain# HG311; Figure 3C). These cells express a non-cleavable mutant of the LexA repressor (G85D) that binds RecA*, but fails to induce SOS (Figure 4A and B) (Markham et al., 1981).

**Figure 4:**
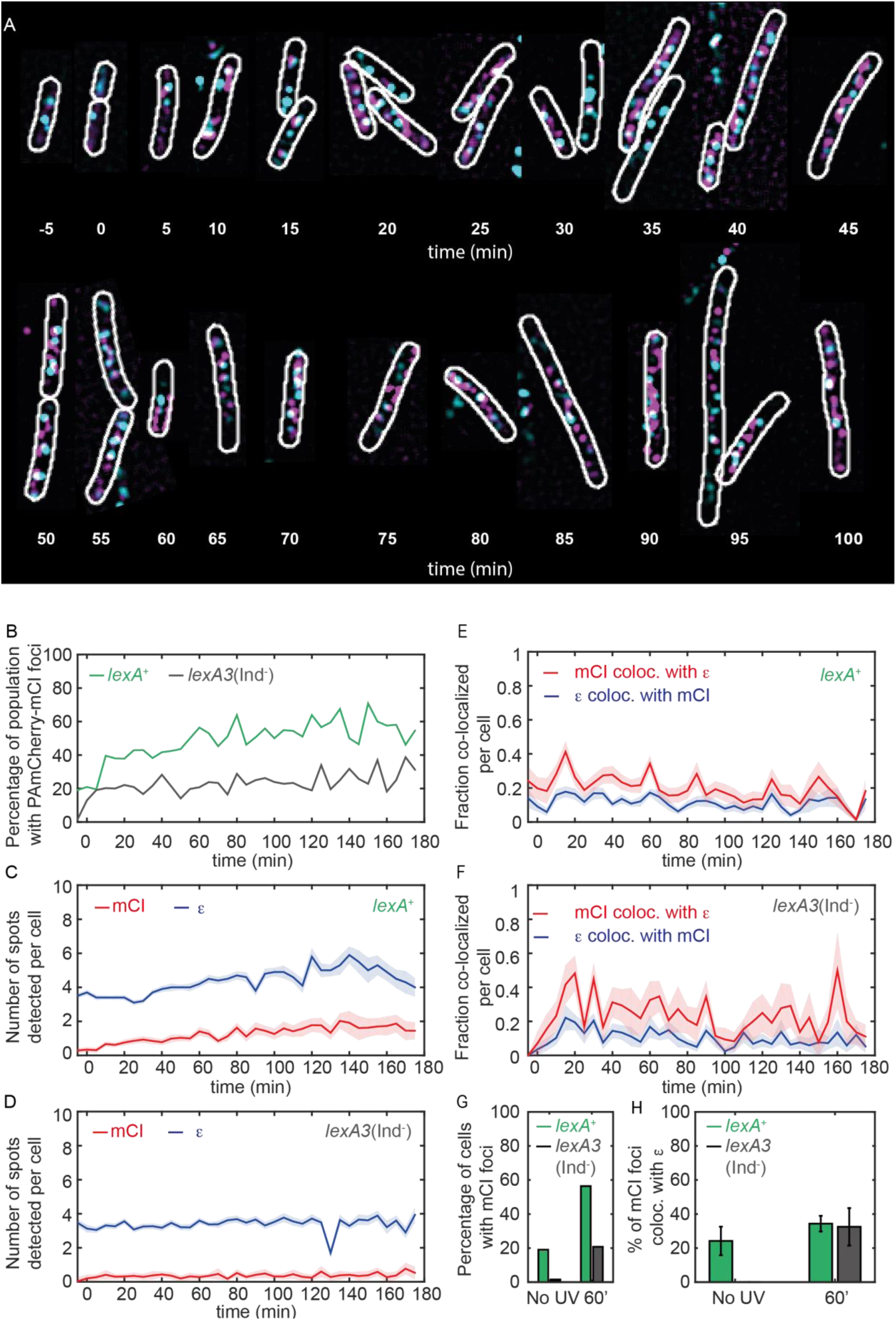
mCI co-localizes with the replisome after UV irradiation. MG1655 cells carrying the ϵ-YPet replisome marker and expressing plasmid PAmCherry-mCI from pBAD-PAmCherry-mCI (strain# HG267) were grown in the presence of 5×10^−4^ % L-arabinose and irradiated with 20 Jm^−2^ of UV-irradiation followed by imaging for three hours. Examples of **A.** *lexA*^+^ (strain# HG267) provided at indicated time points **B.** The percentage of cells imaged at each time point with at least one PAmCherry-mCI focus is shown for *lexA*^+^ (green) and *lexA3*(Ind^-^) (black) cells. Number of replisome foci and PAmCherry foci were counted for each time point per cell for **C**. *lexA*^+^ and **D**. *lexA3*(Ind^-^) cells. In cells exhibiting at least one replisome focus and one PAmCherry-mCI focus, the fraction of replisomes co-localizing with PAmCherry-mCI was determined (blue) and the fraction of PAmCherry-mCI co-localizing with replisomes was determined (red) for **E.** *lexA*^+^ and **F.** *lexA3*(Ind^-^) cells. **G.** Bar plots summarizing percentage of cells exhibiting at least one mCI focus for *lexA*^+^ (green) and *lexA3*(Ind^-^) (gray) cells before UV and at 60 min after UV irradiation. **H.** Bar plots summarizing extent of co-localization of ϵ-YPet and PAmCherry-mCI in cells with at least one mCI and ϵ focus. Data are presented as mean ± SEM. 25-150 cells were analyzed for each time point from at least three repeats of each experiment. See also SI Figure 3.

SOS induction resulted in an increase in mCI/RecA foci. In the absence of DNA damage, 19% of all *lexA*^+^ cells exhibited at least one PAmCherry-mCI focus (Figure 4A and 4C (green line), *t* = −5 min; summarized in Figure 4H). In response to UV damage inflicted at *t* = 0 min, we detected an increase in the number of *lexA*^+^ cells with at least one PAmCherry focus to 56% of the population at *t* = 60 min after UV (green line, Figure 4C, total numbers for each time point are presented in SI Figure 3C). The number of replisome foci detected per cell was found to remain relatively constant ranging from 3.5 ± 0.14 (mean ± standard error) in the absence of UV to 4.2 ± 0.2 per cell at *t* = 60 min after UV (blue line, Figure 4D). The number of PAmCherry-mCI foci marking sites of RecA* was found to increase approximately five-fold from 0.3 ± 0.15 per cell before UV irradiation to 1.4 ± 0.25 per cell at *t* = 60 min (red line, Figure 4D). Among cells exhibiting at least one focus, 24 ± 8% (mean ± standard error) of PAmCherry-mCI foci co-localized with replisomes before UV irradiation, and this number increased to 34 ± 5% at 60 min after UV (Figure 4F and 4I).

Blocking SOS induction also limited RecA/mCI focus formation. Only 1.5% of the *lexA3*(Ind^-^) (gray line Figure 4C) population exhibited at least one PAmCherry-mCI focus (compared to 19% for wild-type) (summarized in Figure 4H) in the absence of UV. Additionally, the number of PAmCherry-mCI foci detected remained consistently low, with 0.28 ± 0.17 foci per cell compared to *lexA*^+^ cells (compare red lines in Figure 4E and 4D), suggesting that the inability to cleave LexA results in an absence of available binding sites for PAmCherry-mCI. Strikingly, *lexA3*(Ind^-^) cells did not exhibit PAmCherry-mCI foci that co-localized with replisomes before UV irradiation (Figure 4G). At 60 minutes after UV irradiation, only 21% of the population exhibited PAmCherry-mCI foci compared to 56% in case of the wild-type (Figure 4C, 4H). Of these, 32 ± 11% of the PAmCherry-mCI foci co-localized with replisomes at 60 minutes (Figure 4G, 4I). These results are consistent with the model that some RecA* filaments are formed in cells in the vicinity of replisomes when cells are exposed to UV light. Most RecA* filaments are formed at locations distal to the replisome.

14 ± 5% of the replisomes exhibited co-localization with mCI in *lexA*^+^ cells exhibiting PAmCherry-mCI foci (Figure 4F). These results suggest that in approximately twenty percent of the population RecA filaments are formed during normal growth (Figure 4C, ‘No UV’ time point). However only approximately 24% of these are associated with replisomes (blue line, Figure 4F, ‘No UV’ time point). These RecA* filaments formed at sites of replisomes in the absence of UV light might reflect replication forks engaged in recombination-dependent DNA restart pathways or replication forks stalled at sites of bulky endogenous DNA damage. In contrast, fewer *lexA3*(Ind^-^) cells exhibited PAmCherry-mCI foci suggesting that the non-cleavable LexA(G85D) protein competes with mCI for the same substrates *in vivo*. These results reinforce our observations that mCI and LexA compete for the same binding substrates *in vivo.*

### RecA forms bundles that are stained by mCI in cells

Previous studies have noted the formation of large macromolecular assemblies of RecA in response to double-strand breaks in cells (Lesterlin et al., 2014). These aggregates of RecA-GFP have been termed ‘bundles’ (Figure 1C and references (Lesterlin et al., 2014; Rajendram et al., 2015)). Our experiments with MG1655/pBAD-PAmCherry-mcI also revealed localizations of mCI that resembled RecA bundles (Figure 5A). These observations were also reproduced in cells expressing YPet-mCI from a pBAD plasmid (strain# HG143). Cells were induced with 10^−3^ % L-arabinose, immobilized in flow cells, irradiated with a dose of 20 Jm^−2^ of UV, and imaged using a time-lapse acquisition protocol. In this experiment, YPet-mCI initially exhibits cytosolic localization at the start of the experiment (Figure 5B, *t* = −5 min (‘No UV’ time point)) and reveals foci and bundles at later time points (Figure 5B, *t* = 1 h and 2 h).

**Figure 5:**
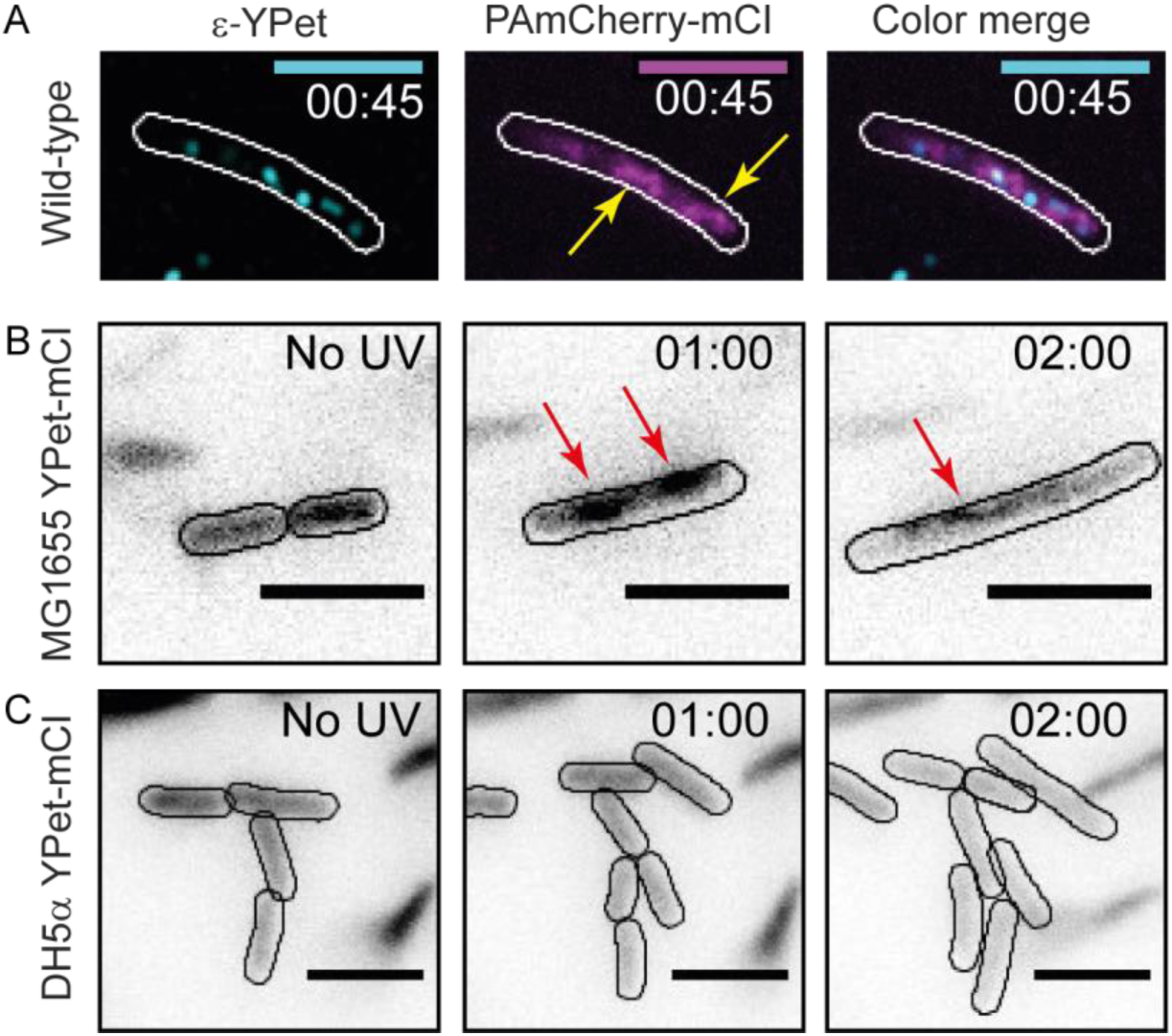
mCI stains RecA bundles after UV-damage. **A.** At late time points in the DNA damage response, PAmCherry-mCI forms large bundles in *recA*^+^ cells. Shown here is an example of an overlay of the mCI signal (magenta) and replisomal ϵ foci (cyan) at *t* = 45 min after 20 Jm^−2^ UV. Yellow arrows point to RecA bundles. For purposes of illustration, peaks in the 514-nm ϵ channel were enhanced using a discoidal average filter. **B.** YPet-mCI also forms bundles (indicated by red arrows) in response to UV-damage in *recA*^+^ cells. **C.** DH5α carrying the *recA1* allele does not exhibit foci or bundle formation upon UV-irradiation under identical conditions as in panel B. Scale bar corresponds to 5 μm. See also SI Figure 4. Cell outlines provided as a guide to the eye.

Next, we tested whether the formation of these bundles required RecA that has wild-type functions. To that end, we imaged YPet-mCI in DH5α cells that carry the *recA1* allele (an inactive mutant, G160D) (Bryant, 1988). These cells did not exhibit foci or bundles that bound to YPet-mCI after UV (Figure 5C, strain# HG242). These data demonstrate that mCI recognizes a specific configuration of wild-type RecA on ssDNA – one that is able to co-operatively bind and hydrolyze ATP.

The UvrD helicase performs a critical role in disassembling RecA filaments in cells (Centore and Sandler, 2007; Lestini and Michel, 2007; Petrova et al., 2015; Veaute et al., 2005). Since mCI stabilizes RecA-ssDNA filaments *in vitro*, we hypothesized that persistent RecA* filaments would lead to constitutive SOS in Δ*uvrD* cells lacking the ability to disassemble RecA*. To test this hypothesis, we imaged Δ*uvrD* cells (strain# HG235) expressing plasmid-based YPet-mCI and found that these cells indeed exhibited constitutive RecA bundles. The presence of constitutive SOS in these cells is further confirmed by the observation of a strong filamentation, even in the absence of any external DNA damage (SI Figure 4).

Taken together, these results demonstrate that: 1) RecA bundles are not only formed by RecA-GFP but also by wild-type RecA during the SOS response and 2) The ability to form a high-affinity complex on ssDNA is critical for the formation of RecA bundles. Further, the lack of mCI features in the *recA1* background suggest that far from being DNA-free aggregates of RecA, these bundles contain an ordered assembly of RecA that is bound to DNA.

### RecA forms storage structures that are not stained by mCI

We then turned our attention to focus on how RecA is stored in cells. Storage structures of RecA would need to satisfy two criteria to be distinguished from complexes active in DNA repair and from polar aggregates representing mis-folded proteins: 1) the size or number of these structures should be proportional to the amount of RecA present in the cell and 2) RecA stored in these structures should be available for biological function when required, that is, after DNA damage. To detect storage structures of RecA in live cells, we imaged *recA-gfp* cells in the absence of DNA damage (strain# HG195). Cells exhibited punctate foci that appear to be positioned outside the nucleoid (Figure 6A). Since the cells did not exhibit additional markers of distress, namely cell filamentation, we presumed that the RecA features correspond to storage structures of RecA. We hypothesized that the size of these punctate foci is dependent on the copy number of RecA in cells. To test this hypothesis, we pursued a strategy involving over-expression of unlabeled RecA from a plasmid in *recA-gfp* cells and measuring whether the foci become larger as the amount of untagged RecA increases and integrates into the structures with the labelled RecA.

**Figure 6:**
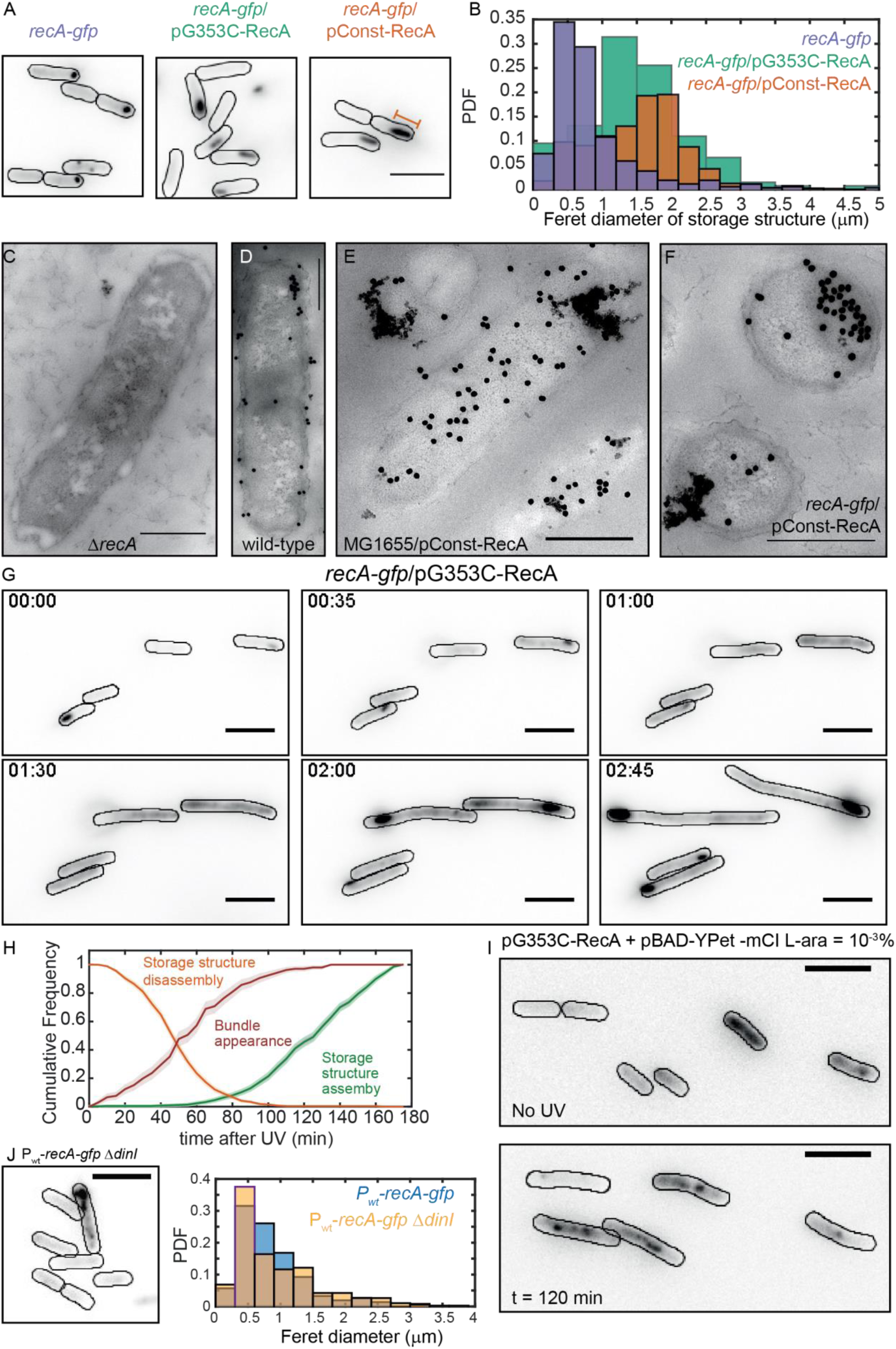
Excess RecA is stored in storage-structures. **A.** Montage of *recA-gfp* (strain# HG195), *recA-gfp*/pG353C-RecA (strain# HG406) and *recA-gfp*/pConst-RecA cells (strain# HG411) imaged in the absence of UV damage. See also SI Movie 3. **B.** Probability density functions of maximum Feret diameter of storage structures in *recA-gfp* (purple), *recA-gfp*/pG353C-RecA (green) and *recA-gfp*/pConst-RecA (orange) cells. Orange bar in panel A represents the maximum Feret diameter for that particular storage structure. Electron microscopy images of **C**. Δ*recA* **D**. wild-type *recA* **E**. *recA-gfp*/pConst-RecA and **F**. MG1655/pConst-RecA cells stained with gold nanoparticles labelled with RecA antibody. Note the appearance of aggregates of gold nanoparticles at locations consistent with those observed in panel A for *recA-gfp*/pConst-RecA cells (panel E). Untagged, over-expressed RecA (F) reveals gold nanoparticle localizations consistent with those expected from RecA storage structures. Scale bar corresponds to 1 μm. **G.** Montage of frames from a time-lapse experiment of *recA-gfp*/pG353C-RecA cells exposed to UV (see also SI Movies 3 and 4). RecA forms storage structures in the absence of DNA damage (0 min) in cells. Storage structures dynamically dissolve after DNA damage (1 h). Storage structures reform by sequestering excess RecA synthesized during SOS after repair (2 h and 2 h 45 min time points). **H.** Cumulative probability distributions of time of solubilization of storage structure (yellow) and time of appearance (light green) of storage structures from *recA-gfp*/pG353C-RecA (strain# HG406) cells. Red line represents cumulative probability distribution of time of first incidence of RecA bundles in *recA-gfp* cells (strain# HG195). Shaded error bars represent standard deviation of the bootstrap distribution obtained by sampling 80% of the data 1,000 times. In each case, 100-150 cells were analyzed that were present for the duration of observation (3 h). Scale bar represents 5 μm. **I.** YPet-mCI does not stain storage structures in MG1655/pG353C-RecA pBAD-Ypet-mCI cells (strain# HG446) in the absence of DNA damage, but forms features after UV damage (shown here is a still at 120 min). Cell outlines provided as a guide to the eye. See also SI Movie 5, and SI Figure 5. **J.** MG1655 cells carrying the *recA-gfp* fusion under the native *recA* promoter and Δ*dinI* (strain# EAW 767) exhibit fewer, and smaller storage structures than *dinI*^+^ cells.

To that end, we created plasmids that expressed wild type RecA protein at two different levels. First, a low-copy plasmid expressed *recA* from the constitutive *recAo281* operator (pConst-RecA; see SI for details) (Uhlin et al., 1982; Volkert et al., 1976). A second version of that plasmid (pG353C-RecA) cut expression in half by incorporating an altered ribosome binding site (RBS) (see SI Figure 5A). We then imaged *recA-gfp* cells carrying one or the other of these plasmids. Time-lapsed imaging of undamaged cells (5 min intervals for 3h) revealed that the RecA-GFP signal was confined to a single large feature (Figure 6A). To quantify the size of RecA features in the absence of DNA damage, we measured the maximum Feret diameter (referring to the largest physical dimension of the structure; Figure 6A) of the feature at a threshold above the background (Figure 6B, see SI for details). Comparison of the Feret diameters of features in *recA-gfp* cells carrying the pConst-RecA and pG353C-RecA vectors revealed a strong dependence on the expression level of untagged wild-type RecA protein (Figure 6B). *recA-gfp*/pConst-RecA cells exhibited larger features than *recA-gfp*/pG353C-RecA cells or *recA-gfp* cells alone. Notably, the storage structures exhibited cross-sections that were circular (in the case of *recA-gfp cells)* or elliptical (in the case of *recA*-gfp/pConst-RecA or *recA-gfp/*pG353C-RecA cells) unlike the previously described thread-like filamentous RecA-bundles (Kidane and Graumann, 2005; Lesterlin et al., 2014). In the absence of DNA damage, these structures were stably maintained in cells in *recA-gfp*/pG353c-RecA cells (SI movie 3). Upon cell division, these structures were disproportionately inherited by daughter cells. Notably, cells did not exhibit markers of distress consistent with induction of SOS, suggesting that these storage structures do not impede DNA replication during growth, and are likely not assembled on DNA. Since the presence and size of these features is dependent on the amount of RecA present in cells, henceforth we refer to them as a ‘storage structures’.

To exclude the possibility that RecA-GFP storage structures are artefacts of the GFP fusion, we collected electron-microscopy images of immunogold-stained RecA in four genetic backgrounds: 1) Δ*recA*; 2) wild-type MG1655; 3) MG1655/pConst-RecA; and 4) *recA-gfp*/pConst-RecA (Figure 6 panels C-F, respectively). As expected, clusters of gold-labelled RecA antibodies could be observed in all samples except Δ*recA* cells (Figure 6C). Cells carrying the over-expresser plasmid pConst-RecA exhibited strong RecA staining that was localized to the membrane (Figure 6E, 6F. See SI Figure 5B for additional examples). These results support the conclusion that excess RecA is stored in the form of membrane-associated, storage structures even in cells carrying untagged, wild-type RecA.

Stored RecA should be made available to support repair during the SOS response. We tested this hypothesis by exposing *recA-gfp*/pG353C-RecA cells to ultraviolet radiation and monitoring the dynamics of the storage structures in time-lapsed fashion. We found that the storage structures dissolve within one hour after introducing damage, flooding the cell with RecA-GFP (Figure 6G, *t* = 1 h). At later time points, the storage structures re-appeared at locations close to the poles (Figure 6G, *t* = 2 h 45), suggesting that the RecA is stored away until needed (see also SI Movie 4). We characterized the dynamics of storage structure disassembly by plotting the cumulative probability of loss of storage structures for the population of cells that possessed a distinct storage structure as a function of time (Figure 6H; orange curve). We found that for *recA-gfp*/pG353C-RecA cells, half of the storage structures were lost within 45 ± 5 min after UV damage (see SI for details). We then plotted the cumulative probability distribution of time of appearance of storage structures after SOS induction (Figure 6H, green curve). In these cells, half of the population of storage structures that formed after UV damage, did so after 135 ± 5 minutes after UV.

Next, we investigated whether mCI interacts with these storage structures. Fluorescence imaging of plasmid-based YPet-mCI expressed from the pBAD promoter in wild-type cells expressing pG353C-RecA (strain# HG446) did not reveal any morphological features consistent with those of the storage structures observed in *recA-gfp*/pG353C-RecA cells. YPet-mCI was found to be cytosolic in the absence of DNA damage, suggesting a lack of stable association with RecA storage structures (Figure 6I, ‘No UV’ time point). As noted earlier, foci were rare. As described before, in response to UV irradiation, cytosolic YPet-mCI was found to form foci and bundles (Figure 6I, 120 min time point; see also SI Movie 4). These results suggested that YPet-mCI does not interact with RecA storage structures. Observations of cells expressing plasmid-based mCI in *recA-gfp*/pG353C-RecA revealed no detectable influence of mCI on the morphology of these structures (SI Figure 5C), reinforcing the interpretation that mCI does not interact with storage structures of RecA in the absence of damage.

To further confirm that storage structures are indeed distinct from RecA bundles, we characterized the kinetics of RecA-GFP bundle formation in *recA-gfp* cells (strain# HG195). In these cells, RecA storage structures are small (Figure 6A) and indistinguishable from repair foci. Upon UV irradiation, cytosolic RecA-GFP forms foci that progress to form large, cell-spanning bundles over the course of several tens of minutes (see SI Movie2), unlike RecA storage structures observed in the *recA-gfp*/pG353C-RecA cells. Plotting a cumulative probability distribution of time of incidence of bundle formation revealed that half of all bundles in *recA-gfp* cells appear by 60 minutes after UV (Figure 6H, red curve). This timing is consistent with measurements of incidence of bundle formation during double-strand break repair (Lesterlin et al., 2014).

### DinI promotes the formation of storage structures in cells

The DinI protein is a modulator of RecA function (Lusetti et al., 2004a; Lusetti et al., 2004b; Renzette et al., 2007). In solution, the C-terminal tail of DinI mimics ssDNA, enabling interactions with monomeric RecA (Ramirez et al., 2000). Since free RecA assembles to form storage structures, we next investigated whether storage of RecA was influenced by DinI. Considering that expression level of RecA influences storage structure formation, we first constructed a strain carrying the *recA-gfp* chromosomal fusion under its native wild-type promoter (P_wt_-*recA-gfp*; strain# EAW428). We deleted *dinI* in this background (strain# EAW767). In the absence of DNA damage, we detected storage structures in fewer Δ*dinI* cells (27% of 702 cells) compared to *dinI*^+^ cells (43% of 855 cells). Additionally, these structures were smaller (see Figure 6J). Over-expression of DinI from pBAD-DinI in Δ*dinI* cells further confirmed this result: cells recovered storage structures in the presence of L-arabinose (see SI Figure 5D). These findings suggest that RecA storage structure formation may be promoted by DinI.

## Discussion

In this work, we have used the C-terminal fragment of the λ repressor in conjunction with single-molecule imaging techniques in live cells to examine RecA protein dynamics in response to SOS induction. In the absence of DNA damage, we see that RecA is largely sequestered in storage structures. Upon UV irradiation, these storage structures dissolve and the cytosolic pool of RecA rapidly nucleates to form early SOS signaling complexes, followed by RecA bundle formation at later time points. Our analysis indicates that the bundles are bound to DNA and may be single extended RecA nucleoprotein filaments. Upon completion of repair, RecA storage structures reform. Our use of the mCI reagent, which associates with DNA-bound and activated RecA* complexes, allows us to eliminate the ambiguity associated with earlier observations utilizing RecA fusion proteins with limited functionality and for the first time provide access to the spatial and temporal behavior of the various forms of RecA structures within the cell. In addition, whereas some RecA foci that form after DNA damage co-localize with replisomes, the majority do not.

We set out to use binding partners of RecA to probe intracellular localization of SOS-signaling RecA complexes. Several proteins associated with the SOS response, notably LexA, UmuD, and the λ repressor, interact with RecA* to effect autocatalytic cleavage. The interaction of these proteins with the activated RecA nucleoprotein filament has been a subject of intense investigation (Cohen et al., 1981; Gimble and Sauer, 1985; Little, 1982). Even though each of these proteins interacts with a different set of residues on the RecA* filament, they all occupy the helical groove of the RecA filament prior to auto-proteolysis (Frank et al., 2000; Galkin et al., 2009; Yu and Egelman, 1993). We sought to exploit this key feature by using a fluorescently labelled C-terminal fragment of λ repressor CI (denoted mCI) to visualize RecA-DNA complexes. The mCI construct binds specifically to RecA* (Figure 2, SI Figure1). Binding of mCI stabilizes RecA* in the ‘active’ conformation capable of mediating transcription-factor cleavage, exhibiting an equilibrium dissociation constant of 36 ± 10 nM and a Hill coefficient of 2.4 ± 0.2 for the binding of mCI to ssDNA-RecA filaments assembled on a dT_40_ ssDNA overhang. Based on previous findings that one mCI contacts two RecA monomers, we estimate that up to six mCI molecules can decorate the RecA-ssDNA filament composed of up to 13 RecA monomers on the dT_40_ DNA substrate under conditions of saturating mCI concentration (Galkin et al., 2009; Ndjonka and Bell, 2006). The non-cleavable mCI thus can decorate RecA filaments assembled on DNA. We confirmed that mCI interacts with RecA filaments in live cells by probing SOS induction after UV damage (Figure 3). We found that mCI has the potential to robustly inhibit SOS induction at high concentrations. SOS induction is retained, albeit delayed, at mCI concentrations employed in this study.

The leading model for SOS induction is that replication forks fail at sites of lesions and produce large tracts of ssDNA that templates nucleation of RecA filaments. Visualizing this model in cells has been challenging due to the difficulties associated with co-localization of a high-abundance protein (RecA) with a handful of replisomes. Our strategy involving fluorescently tagged mCI enabled us to examine the location of RecA* foci in nucleoids relative to the replisomes for the first time in *E coli*. We found that the average number of replisome foci did not change after DNA damage, confirming that most replisomes are not disassembled after UV (Figure 4). Live-cell PALM imaging of mCI revealed foci that depended on the presence of wild-type RecA and DNA damage. Surprisingly, 20% of wild-type cells exhibited RecA* foci during normal growth. However, only 24% of these co-localized with replisomes. The remaining 76% of sites of RecA* detected during normal growth did not co-localize with replisomes. Upon exposure to ultraviolet light, 56% of cells exhibited RecA* foci that were visualized by mCI, with up to 35% of the RecA* foci co-localized with replisomes at 60 min in rich media (Figure 4). A previous report on co-localization of RecA-GFP with DnaX-mCherry in *Bacillus subtilis* growing in minimal media reported a basal co-localization of 74.8 ± 8.4 % with an increase to 84.3 ± 5.8 % at 5 min after 40 Jm^−2^ UV treatment (Lenhart et al., 2014). The extent of co-localization of RecA* and replisomes detected in our experiments, in *E coli* cells growing in media that supports multi-fork replication is lower both before and after UV irradiation. The RecA* foci that co-localize with replisomes are likely associated with replisomes that are stalled at sites of DNA damage. We postulate that RecA* foci that are not co-localizing with replisomes are forming in DNA gaps that are formed and left behind by the replisome (Howard-Flanders et al., 1968; Rupp and Howard-Flanders, 1968; Yeeles and Marians, 2013). Notably, most foci of the translesion DNA polymerases IV and V also form at nucleoid locations that are distal from replisomes, both before and after SOS induction (Henrikus et al., 2018; Robinson et al., 2015).

The large cell-spanning structures termed RecA threads or bundles (we have adopted the latter term) (Kidane and Graumann, 2005; Lesterlin et al., 2014; Rajendram et al., 2015) that form after SOS induction deserve special mention. Following the initial phase of RecA* formation, cells expressing YPet-mCI, PAmCherry-mCI or RecA-GFP exhibited large RecA bundles. The formation of these bundles was also contingent upon the presence of wild-type RecA. The *recA1* allele in DH5α failed to support focus or bundle formation, consistent with the inability of the RecA(G160D) in DH5α to induce SOS and HR functions (Bryant, 1988). Additionally, cells lacking UvrD exhibited constitutive RecA bundles. These bundles are thus a hallmark of the DNA damage response and may have special functionality in the homology search required for recombinational DNA repair (Lesterlin et al., 2014). Here, we show that the bundles bind to our mCI probe. This implies that the bundles are either bound to DNA and thus activated as RecA*, or at a minimum are in a RecA*-like conformation that permits mCI binding. Interestingly, despite the differences in the nature of the DNA damage inflicted, the timing of RecA bundle formation in our UV experiments coincided closely with that of RecA bundles observed upon induction of site specific double-strand breaks in the chromosome (Lesterlin et al., 2014). The bundles may thus be nucleated by RecA binding to either gaps or resected double strand breaks. At later time points, polymerization of RecA filaments nucleated at ssDNA gaps could extend onto dsDNA, and manifest as bundles in our experiments. RecA bundles also interact with anionic phospholipids in the inner membrane (Rajendram et al., 2015). Notably, UmuC also localizes primarily at the inner-membrane upon production and access to the nucleoid is regulated by the RecA* mediated UmuD cleavage (Robinson et al., 2015). This transition occurs late in the SOS response (after 90 min), at a time-point when most, if not all of the cells in the population exhibit RecA bundles. The origins, maturation and additional catalytic roles of RecA bundles in the SOS response require additional investigation.

Our experiments enable us to distinguish storage structures from SOS signaling complexes and RecA bundles based on three qualities: 1. storage structures dissolve after UV damage whereas RecA bundles are formed in response to DNA damage. 2. storage structures often exhibit a polar localization, whereas RecA-bundles form along the cell length. 3. the SOS signaling complexes and RecA bundles are visualized by binding to mCI, whereas the RecA storage structures are not. Finally, we found that DinI promotes storage structure formation: cytosolic RecA in normal growing cells was found to be sequestered in structures by simply over-expressing DinI from a plasmid.

Taken together, these data for the first time provide a full picture of a process that was first hypothesized by Story and co-workers in 1992 suggesting that RecA can undergo a phase transition to form DNA-free assemblies in live cells and redistribute into the cytosol where it becomes available for DNA-repair functions (Figure 7). Within a few minutes after encountering bulky lesions, replication forks synthesize ssDNA substrates that are rapidly coated by cytosolic RecA to form RecA*. These RecA* enable auto-proteolysis of LexA to initiate the SOS response and increase the levels of cellular RecA protein. Meanwhile, storage structures of RecA dissolve, making RecA available for biological functions. The RecA* foci elongate over several hours into elaborate bundles that may have multiple functions. Finally, excess RecA is sequestered away into storage structures approximately two hours after DNA damage, after DNA repair is complete and normal growth is restored.

**Figure 7:**
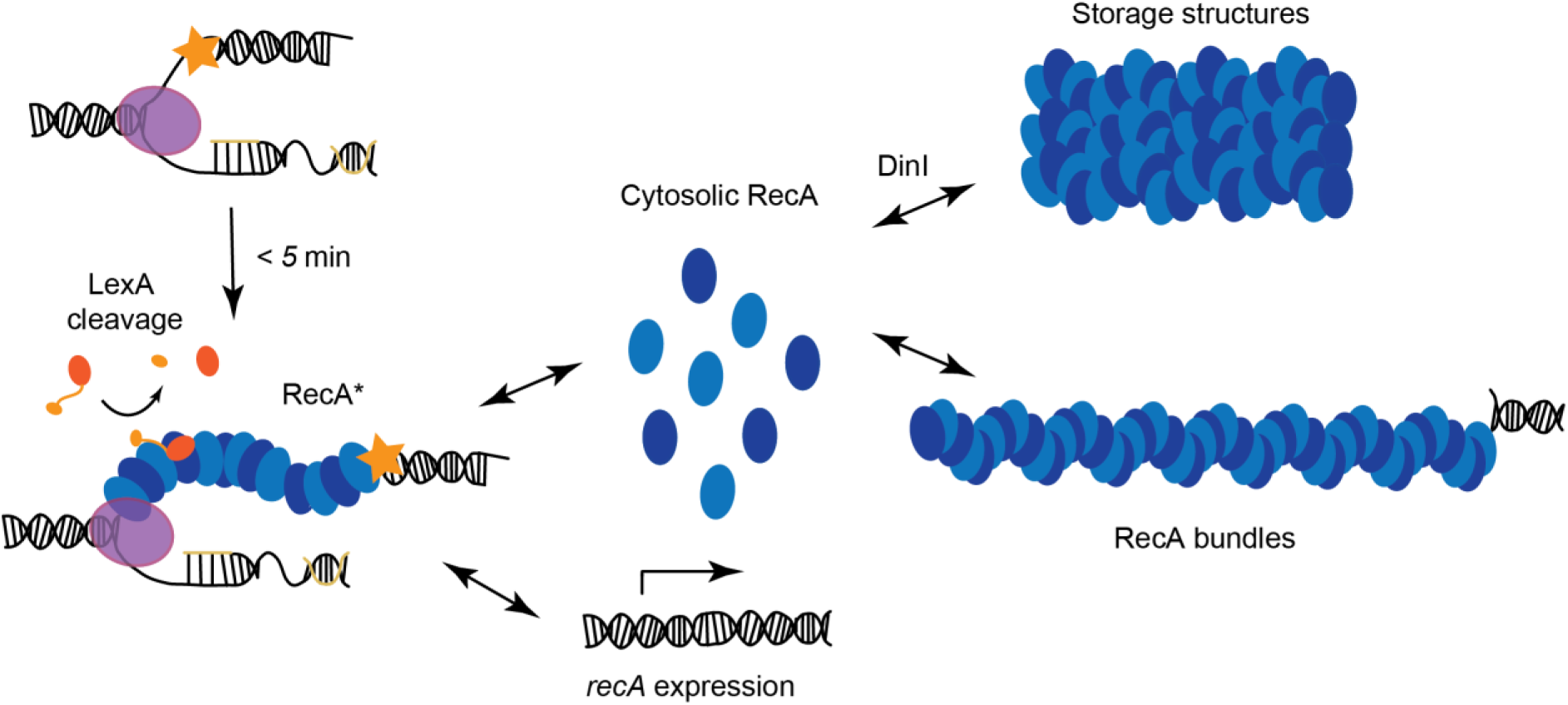
Model for organization of RecA complexes after DNA damage. Model for storage and re-distribution of RecA after DNA damage. Detection of UV damage leads to formation of ssDNA-RecA filaments at sites of replisomes. These ssDNA-RecA filaments catalyze auto-proteolysis of LexA to induce SOS and upregulate expression of RecA. Storage structures of RecA dissolve in response to DNA damage to make RecA available for repair and recombination.

## Author contributions

Conceptualization: H.G, M.M.C, and A.M.V.O.; Methodology: H.G. and A.M.V.O.; Formal Analysis and Software: H.G. and B.P., Investigation, H.G., B.P., J.L. S.J. and R.W.; Writing – Original Draft: H.G.; Writing – Review & Editing: H.G., M.M.C., R.W., and A.M.V.O.; Funding Acquisition: A.M.V.O.; Resources: E.A.W., R.W.; Supervision: A.M.V.O.

## Acknowledgements

We thank Douglas Weibel for the MG1655 RecA-GFP strain. We thank the Alon lab for the SOS-reporter plasmids. We thank Amy McGrath and Celine Kelso for technical assistance with ESI-MS analyses of purified proteins. R.W. was supported by the NICHD/NIH Intramural Research Program. M.M.C is supported by NIH Grant GM32335. A.M.V.O. acknowledges support by the Australian Research Council (DP150100956 and FL140100027).

